# NEU1 and NEU3 enzymes alter CD22 organization on B cells

**DOI:** 10.1101/2021.04.29.441886

**Authors:** Hanh-Thuc Ton Tran, Caishun Li, Radhika Chakraberty, Christopher W. Cairo

## Abstract

The B cell membrane expresses sialic acid binding Immunoglobulin-like lectins, also called Siglecs, that are important for modulating immune response. Siglecs have interactions with sialoglycoproteins found on the same membrane (cis ligands) that result in homotypic and heterotypic receptor clusters. The regulation and organization of these clusters, and their effect on cell activation, is not clearly understood. We investigated the role of human neuraminidase enzymes, NEU1 and NEU3, on the clustering of CD22 on B cells using confocal microscopy. We observed that native NEU1 and NEU3 activity influence the cluster size of CD22. Using single-particle tracking, we observed that NEU3 activity increased the lateral mobility of CD22, which was in contrast to the effect of exogenous bacterial NEU enzymes. Moreover, we show that native NEU1 and NEU3 activity influenced cellular Ca^2+^levels, supporting a role for these enzymes in regulating B cell activation. Our results establish a role for native NEU activity in modulating CD22 organization and function on B cells.

## INTRODUCTION

B cell receptors (BCRs) are responsible for antigen recognition leading to B cell activation and proliferation in immune response. Regulation of B cell activation involves co-receptors that fine-tune BCR signaling. One widely studied negative regulator of BCR is CD22 (Siglec-2), a member of the sialic-acid binding Immunoglobulin-like lectin (Siglec) family.(1) The structure and organization of CD22 on the cell membrane plays a crucial role in its activity. CD22 is a transmembrane protein containing seven Immunoglobulin domains (Ig) that adopts a rod-like structure; the N-terminal Ig domain specifically recognizes terminal α2,6-sialic acids.(2) The cytoplasmic portion of CD22 contains Immunoreceptor Tyrosine Inhibitory Motifs (ITIMs) that dampen cellular response.(3) CD22’s lectin domain can interact with sialosides from cis- or trans-ligands; however, the high density of cis sialosides on the membrane results in the formation of hetero- and homotypic clusters of CD22. The N-link glycans of CD22 allow for homotypic interactions,(4,5) while heterotypic cis-binding partners, including CD45,(6,7) are known. Although CD22’s ligand binding is somewhat weak,(8) high-affinity trans-ligands can overcome cis-interactions despite their prevalence.(9,10) High-affinity multivalent displays of CD22 ligands including liposomes,(11,12) polymers,(4,10,13) and synthetic scaffolds(14) have been proposed as modulators of B cell activation.

The complexity of CD22 interactions and organization on the B cell membrane remains an active area of investigation. Previous studies have observed nanoclusters of CD22, with a minor role for the actin cytoskeleton in lateral mobility.(15) Despite the large number of sialylated glycoproteins on the membrane, only a limited number of these have been identified as *in situ* CD22 ligands.(5) This observation may indicate a role for membrane microdomains in enforcing specific interactions, as CD45 and IgM have close associations with lipid rafts.(16–19) A common strategy for investigating the role of CD22-sialoside interactions is to use sialic acid-cleaving enzymes: *neuraminidases* (NEUs, also called *sialidases*).(20) Recombinant NEU from bacteria have been used for this purpose as research tools. NEU from *Clostridium perfringens* (NanI) has substrate preference for α2,3-glycoproteins, while the sialidase of *Athrobacter ureafaciens* (siaAU) has a broader range of substrate specificity cleaving α2,3-, α2,6-, or α2,8-linked gangliosides or glycoproteins. Exogenous NEU reagents have helped establish the importance of CD22-sialoside interactions; however, there has been very little work to investigate the role of native NEU enzymes on this system. There are four human NEU isoenzymes (hNEU): NEU1, NEU2, NEU3, and NEU4 and they have important cellular functions and roles in health and disease,(21) including atherosclerosis,(22) malignancy,(23–25) and neurodegenerative diseases.(26–29) Together with glycosyl transferase enzymes (GTs), NEU regulate sialic acid content in cells.(20) Thus, native NEU activity could act as a regulator of CD22 organization, B cell activation, and immune response.

Here, we investigated the influence of the cytoskeleton and native NEU activity on the membrane organization and dynamics of CD22. By utilizing confocal microscopy and single-particle tracking, we found that clustering and diffusion of these receptors are dependant on both cytoskeletal structure and changes in glycosylation of B cells. We confirmed our findings by performing knockdown of hNEU expression in model B cells. Finally, we confirmed that native hNEU activity affects B cell calcium levels. We conclude that organization and diffusion of CD22 receptors is dependent on an intact cytoskeleton and the homeostasis of native sialoside ligands.

## MATERIALS AND METHODS

### Confocal microscopy

Cells were grown in R10 media and kept in a humidified incubator at 37 °C and 5% CO2. For confocal microscopy experiments, Raji cells (1 × 10^6^) were counted, centrifuged, and re-suspended in Hank’s Balanced Salt Solution (HBSS). The cells were washed and treated with NanI (*Clostridium perfringens*), siaAU (*Athrobacter ureafaciens*), or NEU3 in HBSS, or CytoD or LatA in HBSS with 0.005 % DMSO at 37 °C for 1 hour, then fixed using 1% PFA on ice for 30 min. Samples were treated with 1 *μ*L/mL mouse anti-human IgM (clone IM260, Abcam cat# ab200541) or mouse anti-human CD22 (clone HIB22, BD Pharmingen cat# 555423) at 4 °C overnight, and stained with goat anti-mouse IgG (polyclonal, Sigma-Aldrich cat# M8642) conjugated with Alexa Flour 647 (AF647) at room temperature for 1 h. The loading of the fluorophores was approximately 2 dyes/protein. After washing, samples were transferred to 24-well plates (Corning, Inc.) with circular cover glass slides pre-treated with poly-L-lysine (PLL), spun at 300 x g for 15 min, washed, and glass slides were mounted onto microscopy slides with Slowfade Antifade (Thermo Fisher, cat# S2828) and sealed using Cytoseal 60. Samples were imaged on a laser scanning confocal microscope (Olympus IX81 with 60X objectives). Twenty cells from each condition were chosen for analysis based on transmitted and fluorescence images, and each cluster was analyzed using the particle analysis function on ImageJ. The data were plotted using the beanplot plugin in R, and statistics were done using Graphpad Prism. Three runs were performed and analyzed, and one representative run is shown for each condition.

### Single-Particle Tracking

Raji cells were grown in R10 media as previously described.(14) For each condition, cells (1 × 10^6^) were counted and washed with HBSS buffer three times. The cells were then re-suspended in HBSS and treated with NanI (10 mU/mL), siaAU (5 mU/mL or 10 mU/mL), or NEU3 (10 mU/mL). For CytoD and LatA, the cells were re-suspended in 0.005 % DMSO and the compounds were added to a final concentration of 100 mM. For all conditions, the cells were incubated for 1 h at 37 °C and 5% CO2. After incubation, the cells were centrifuged and washed three times with HBSS buffer at 200 x g for 15 min, re-suspended in HBSS and labelled with anti-CD22 conjugated with AF647 (1h at room temperature). After labelling, the cells were washed three times in buffer then transferred to 24 well-plates with circular glass cover pre-treated with PLL. The plate was then spun at 300 x g for 15 min, and the cover glass was mounted to microscope slides with buffer and sealed with Cytoseal 60. To record the trajectories, TIRFM was performed on Nikon Ti microscope. The angle of incidence was set at ∼1100, and videos were taken at 10 FPS for 10 seconds.(30) All the videos were taken within one hour of slide preparation. Trajectories were analyzed using UTrack software in MATLAB, and the coefficient of diffusion was calculated using MATLAB.(30)

### Transfection of Raji cells

Raji cells were grown as described and passaged 24 h prior to transfection.(14) Cells (24 × 106) were washed and re-suspended into 750 µL of electroporation buffer (PBS with no Ca^2+^or Mg^2+^). For each knock-down condition, 15 µL from the 20 µM stock siRNA solution was added, while 15 µL of siRNA buffer was added for the no-treatment control. Cells were mixed by pipet and transferred to a 2 mm electroporation cuvette. Cell samples were left on ice for 20 min, and electric shock was applied using a Bio-Rad electroporator (0.6 kV, 50 µF, and 350 Ω). The cuvettes were then immediately transferred and left on ice for 30 min. Transfected cells were transferred to T25 cell culture flasks and pre-warmed R10 growth medium was added so the final volume for each condition was 6 mL. Cells were kept in a humidified incubator at 37 °C with 5% CO2 for 24 h.

### Ca^2+^ activity assay of Raji cells

For each condition, Raji cells were counted and re-suspended to a final concentration of 1 × 10^6^ cells/mL. Cells were then treated with NEU at a final concentration of 10 mU/mL or with 100 nM neuraminidase inhibitors. For treatments of enzyme with inhibitors, these were incubated together for 30 min before addition to cells. The cells were treated with indicated conditions for 1 h at 37 °C with 5% CO2. For transfected cells, cells were transfected and grown 24 hours prior to Ca^2+^ experiments. Cells were washed three times with PBS and re-suspended at 5 × 10^6^/mL in loading buffer (RPMI-1640 supplemented with 1% FBS, 10 mM HEPES, 1mM MgCl2, 1mM EGTA, and 5% penicillin-streptomycin) and treated with 1.5 µM Indo-1 dye in a 37 °C water bath for 30 min while protected from light. After loading, the cells were washed with loading buffer three times, then re-suspended in running buffer (HBSS supplemented with 1% FCS, 1 mM MgCl2 and 1 mM CaCl2), after which they were stored on ice. Using a Fortessa X10 FACS machine, a plot of violet (379 nm) versus blue (450 nm) was created and voltages were adjusted so that a maximum of 5% of unstimulated cells lie within the violet gate. To measure the amount of Ca^2+^ flow into the cells, 500 µL of previously suspended cells from each condition were transferred to FACS tubes incubated in 37 °C water bath for 2 min before running the samples. Ten seconds after the acquisition was initiated to establish the background, the tube was quickly removed and either PBS (unstimulated condition) or anti-IgM (stimulated condition) was added and vortexed before placing the tube back. The total acquisition time was 3 min for each tube and the number of cells analyzed ranged from 150-400 thousand cells. From these data, the percentage of cells emitting violet light from each condition was normalized to that of control unstimulated cells, and a Student’s t-test was performed with GraphPad Prism.

### Western Blot of Raji cells

Transfected cells were centrifuged and supernatant was discarded. The cells were lysed using 100 µL of HEPA buffer with protease inhibitor on ice for 1 h. The lysate was sonicated and centrifuged at 12000 x g for 20 min. Supernatant was collected and the amount of protein was determined using BCA assay. For each condition, 10 µg of protein was mixed with the same volume of 2X gel running buffer with 5% dithiothreitol, followed by 5 min incubation at 95 °C. After heating the samples, they were then transferred to 11% acrylamide gel and SDS-PAGE was run at 110 V for 70 min. After the gel was completed, it was washed with water, and transferred to a nitrocellulose membrane in 1X blotting buffer at 20V for 2.5 h. Once the transfer was completed, the transfer of proteins was confirmed by soaking the membrane in Ponceau S buffer for 5 min, washing the membrane with water, and checking for presence of bands on the membrane. The membrane was blocked in blocking buffer (5% non-fat milk in 1X TBST) on an orbital shaker overnight at 4 °C. The membrane was then washed three times with TBST for 5 min each time. The membrane was stained using primary antibody in TBST with 1:1000 dilution for 1 h at room temperature. The membrane was washed 5 times with TBST, then stained using secondary antibody conjugated with horse radish peroxidase with 1:10000 dilution in TBST for 1 h at room temperature. The membrane was washed and developed using ECL substrate for western blotting and visualized. Percent knock-down was determined by analyzing the intensity of each band using ImageJ. The samples were normalized to the control, and a Student’s t-test was implemented for analysis.

### Lectin blots

CD22 proteins were purified from Raji cells using murine hybridoma cells expressing anti-CD22 antibodies (α-CD22:4213 (10F4.4.1)). The purified sample was validated by SDS-PAGE and the protein concentration were determined using a BCA assay. From the stock solution, 4.4 µg of protein was diluted to a final volume of 200 µL and enzyme concentration of 10 mU/mL, then the samples were incubated at 37 °C for 3 hours. After treatment of the proteins with enzyme, each sample was mixed with the same volume of 2X PAGE sample buffer with SDS, then 40 µL from each sample was subjected to SDS-PAGE with 10% acrylamide at 110 V for 70 minutes. For each of the samples, three lanes were loaded together with control samples. Gels were then taken out, washed three times with water, and transferred to nitrocellulose membrane. The presence of proteins on the membrane was confirmed using Ponceau S, then they were cut so that a control and an enzyme-treated lane was retained. The membranes were washed with TBST and blocked with blocking buffer (5% non-fat milk in TBST) at 4 °C overnight. Membranes were washed three times with TBST, then treated with 1 µg/mL biotinylated lectins (PNA, SNA, MAL) at room temperature for 1 h. The treated membranes were washed 5 times with TBST and treated with streptavidin-HRP for 1 h at 1:10000 dilution, followed by washing the membranes with TBST. The bands were detected using ECL substrate for western blotting under a visualizer. The analysis of the bands was conducted using ImageJ.

## RESULTS

### Cytoskeletal interactions of BCR

To study the effect of cytoskeletal contacts on BCR, we used Raji cells treated with cytoskeletal disruptors *cytochalasin D* (CytoD) and *latrunculin A* (LatA) as a model B cell system. Microclusters of BCR are known to form when B cells are stimulated due to the association of the constant region Cµ4, followed by phosphorylation that induces cellular response.(31) Several studies have observed cytoskeletal interactions with BCR, and the effect of cytoskeletal disruption was increased BCR cluster size and cell activation.(32,33) Cells were treated, fixed, and imaged by confocal microscopy allowing quantitation of cluster size (**Figure 1**).(14) CytoD and LatA were used to disrupt the cytoskeleton.(34–37) We observed a significant increase in clustering of BCR upon treatment with both inhibitors at all concentrations,(15) and there was significantly increased clustering at lower concentration as compared to higher concentrations (2.5 vs 7.5 µg/mL; p < 0.05).(38)

**Figure 1.**
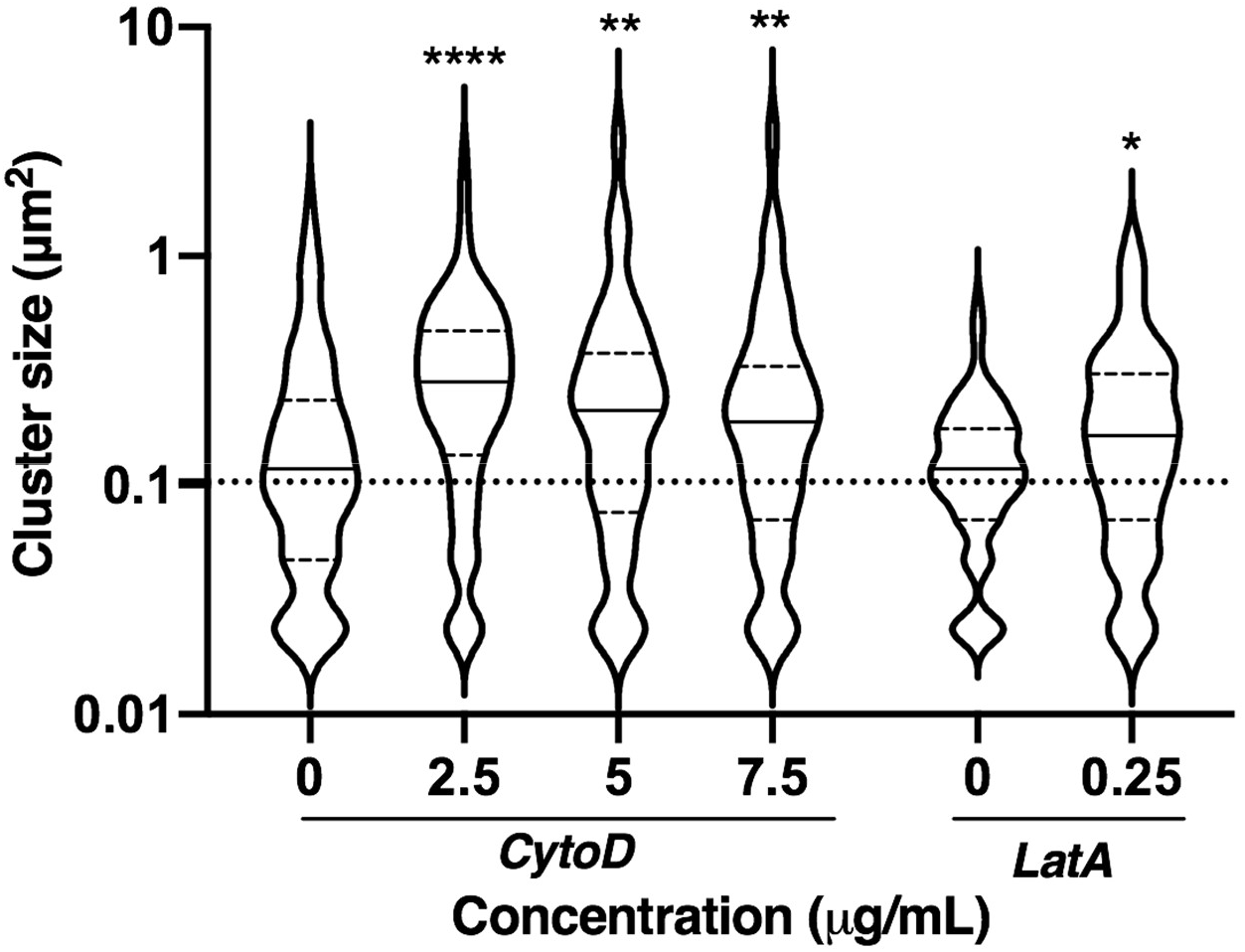
Cluster size of BCR after treatment with cytoskeletal disruptors. Raji cells were treated with cytochalasin D (CytoD) or latrunculin A (LatA) at 37 °C for 30 min. Cells were then fixed and stained with mouse anti-IgM and anti-mouse IgG-AF647 and imaged using confocal microscopy. Data shown are average from 30 cells among 3 biological replicates; cells were analyzed using imageJ and shown as beanplots.(14) Comparisons by student’s t-test are shown relative to respective controls (****, p < 0.0001; **, p < 0.01; *, p < 0.05).

### Cytoskeletal interactions of CD22

We next examined the cytoskeletal interactions of CD22 in a B cell model. Analysis of CD22 clustering after treatment with cytoskeletal disruptors is shown in **Figure 2**. We observed that CD22 cluster size increased significantly when cells were treated with CytoD. Interestingly, the cluster size was increased at intermediate concentrations (2.5 – 7.5 µg/mL); however, this effect was lost at higher concentration (10 µg/mL). Similar trends have been observed in tight junctions, and may be attributed to different populations of actin within the cell.(38,39) Previous work has suggested that CytoD does not alter CD22 clustering or organization in the membrane; however, these studies were performed at higher drug concentrations (10 µM) and our results suggest that lower concentrations may be more appropriate in this system.(15) Treatment of cells with LatA showed no effect on the cluster size of CD22 at the concentrations tested (0.050 – 0.500 µg/mL). One explanation for this difference is the disparate mechanisms of actin disruption used by these two inhibitors.(37) Alternatively, the incubation time with LatA (30 min) may have been insufficient.(40,41)

**Figure 2.**
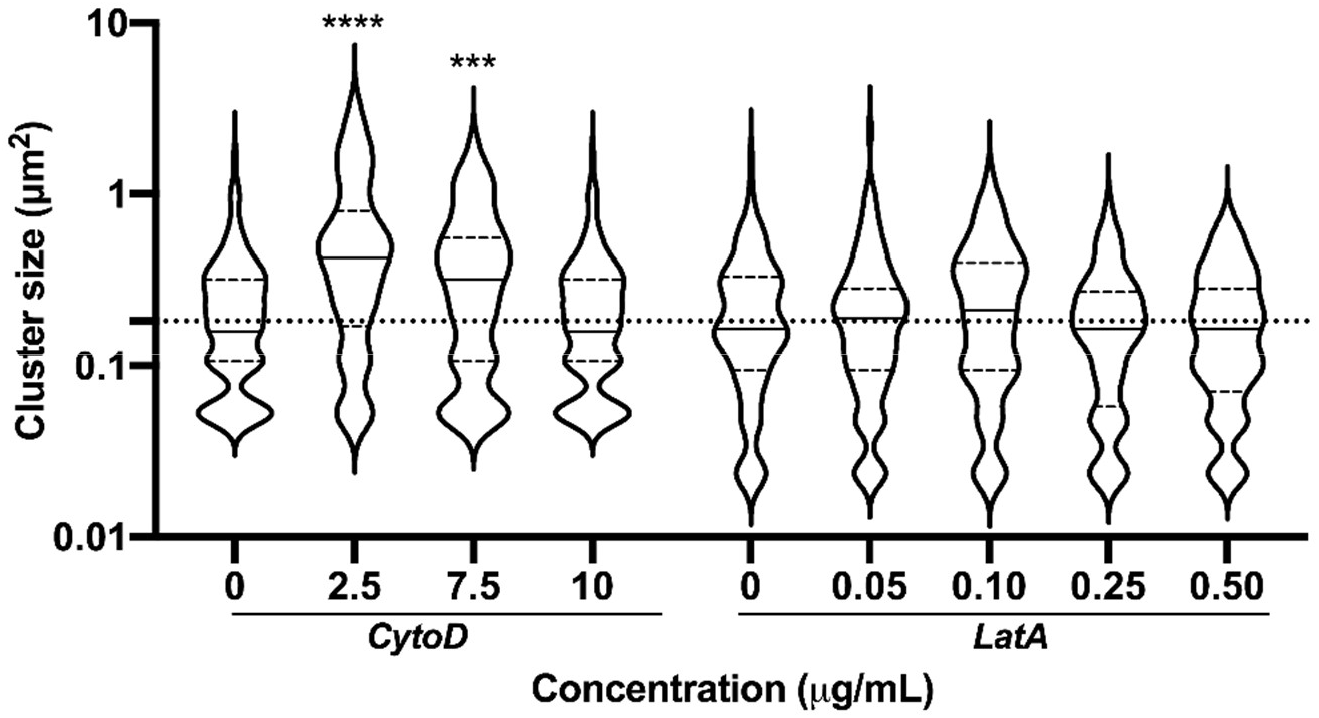
Cluster size of CD22 after treatment with cytoskeletal disruptors. Raji cells were treated with cytochalasin D (CytoD) or latrunculin A (LatA) at 37 °C for 30 min. Cells were then fixed and stained with mouse anti-CD22 and anti-mouse IgG-AF647 and imaged using confocal microscopy. Data shown are average from 30 cells among 3 biological replicates; cells were analyzed using imageJ and shown as beanplots.(14) Comparisons by student’s t-test are shown relative to respective controls (****, p < 0.0001; ***, p < 0.001).

We next investigated if cytoskeletal interactions had a significant influence on the lateral mobility (diffusion) of CD22 in the membrane. We employed single-particle tracking (SPT) using Total Internal Fluorescence Microscopy (TIRFM) to measure changes in CD22 membrane diffusion.(30) Raji cells were stained with minimal amounts of Alexa Flour 647 (AF647) - conjugated primary antibody to CD22. This sparse labelling allowed visualization of CD22 trajectories on live cells, which could be converted to rates of diffusion (**Figure 3**).(42) This method allowed us to compare lateral diffusion of proteins in control and cytoskeleton-disrupted conditions. Treatment with CytoD at low concentration (2.5 µg/mL) significantly decreased CD22 diffusion; however, at higher concentrations (10 µg/mL) this effect was lost. Treatment with LatA (0.25 µg/mL) also showed a significant decrease in lateral mobility of CD22.

**Figure 3.**
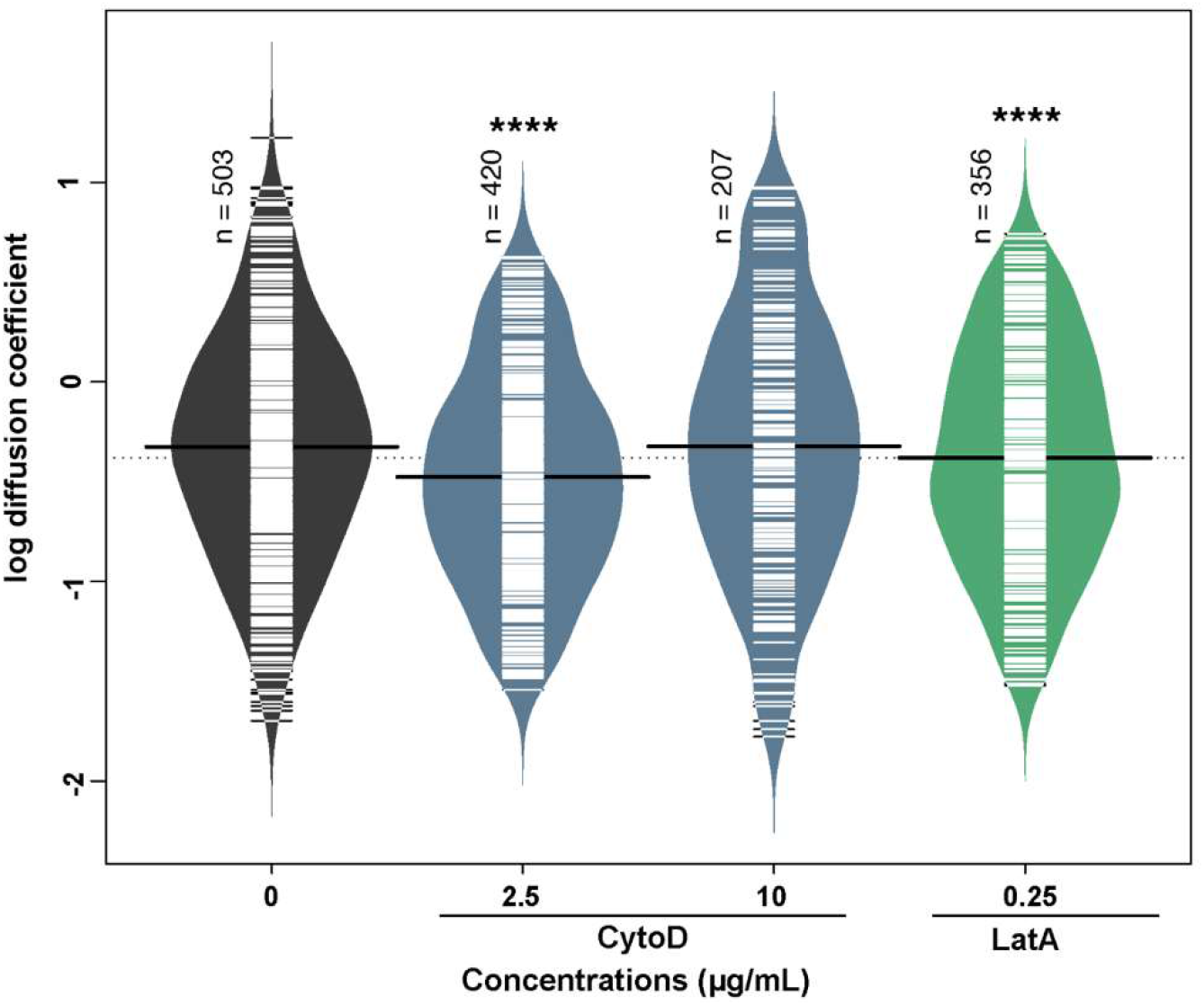
Lateral mobility of CD22 after treatment with cytoskeletal disruptors. Raji cells were treated at 37 °C for 30 min. Lateral mobility was analyzed using single-particle tracking with TIRFM videos recorded at 10 FPS for 10s.(30) Diffusion coefficients are given as log(D), where D is in units of × 10^−10^ [cm^2^s^−1^] or × 10^−2^ [μm^2^s^−1^]. 150 cells among 3 biological replicates were analyzed, and values were compared to control using a student’s t-test (****, p < 0.0001). Beanplots were generated using R software. Individual data points are represented by short white lines, a solid black line indicates the average for each condition, and the dotted line represents an average across all populations.

### NEU1 and NEU3 activity alter CD22 cluster size

Considering that the lectin activity of CD22 is dependent on sialylated cis ligands, we next investigated if native hNEU enzymes could alter CD22 membrane organization. CD22 is found in homotypic clusters,(5,43) and also has cis interactions with other sialylated proteins, including CD45.(44–46) We developed an siRNA knockdown protocol using electroporation for NEU1 and NEU3 enzymes,(47) as lymphocytes are often difficult to transfect using lipid-based methods. The reduced expression of NEU1 and NEU3 was confirmed by western blot of the transfected cells (**Figure 4A, 4B, S1**). We found that B cells treated with siRNA for *Neu1* or *Neu3* had expression of the enzymes reduced by approximately half. Viability of the cells by hemocytometer after treatment showed no significant decrease for *Neu1* siRNA, while *Neu3* siRNA did show a decrease in viability (**Figure S2**). We proceeded to determine if NEU1 and NEU3 knockdown (KD) cells showed evidence of changes to CD22 membrane organization. Analysis of clustering in these cells found a significant increase in CD22 cluster size in NEU1 KD cells while NEU3 KD cells had a significant decrease in cluster size, suggesting these two isoenzymes play different roles in regulating CD22 organization (**Figure 4B**). This observation can be partly attributed to the different substrate specificities of the two enzymes – with NEU1 known to prefer glycoprotein substrates and NEU3 to prefer glycolipids.(20) Additionally, the expression levels of these two enzymes may vary in lymphoid cells, with NEU1 generally being found at higher expression in many cell types.(48,49)

**Figure 4.**
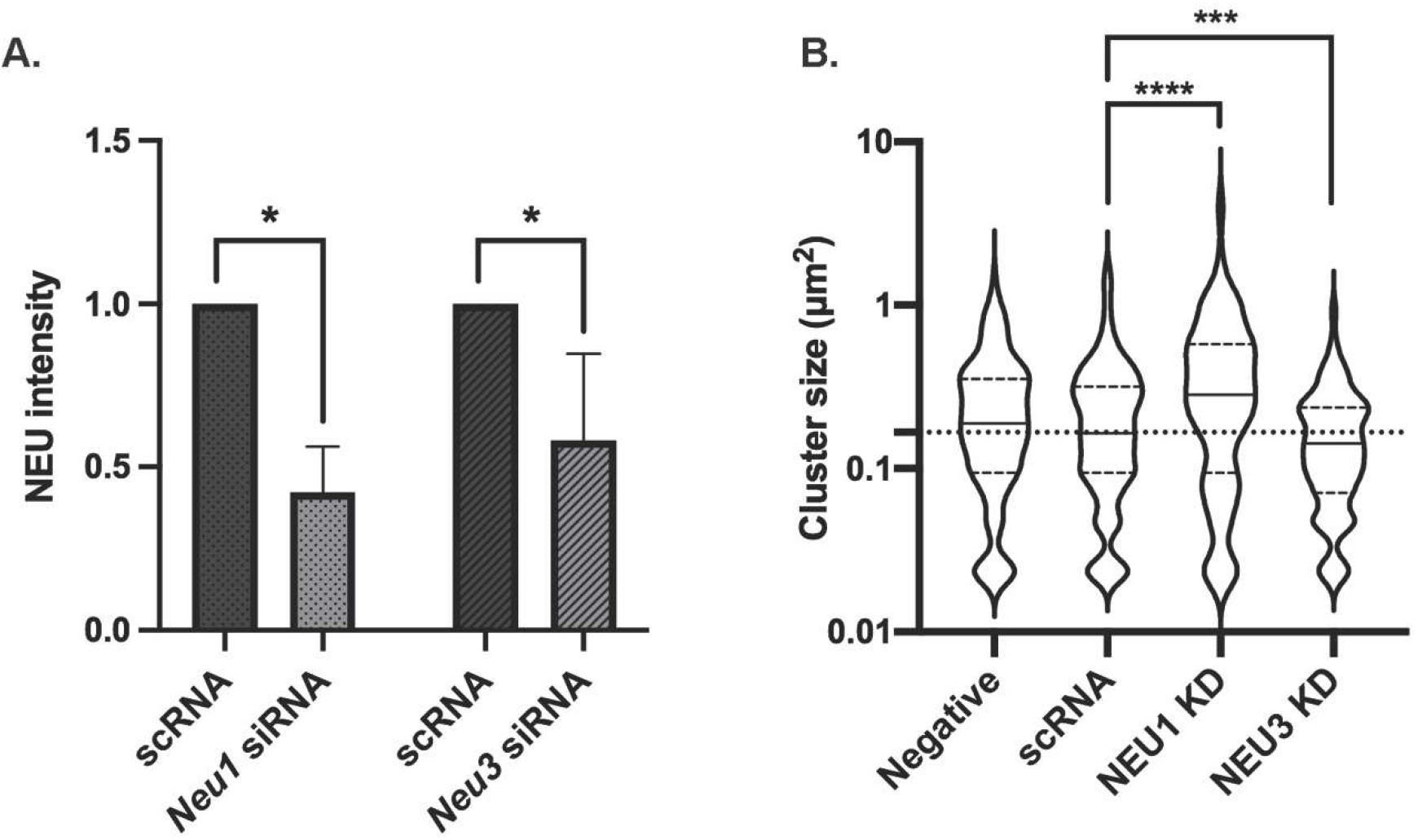
CD22 cluster size is altered by NEU1 and NEU3 knockdown. Raji cells were transfected with siRNA targeting *Neu1* or *Neu3* using electroporation and grown for 24 h. (**A**) Western blots confirmed reduced expression of NEU1 and NEU3. (**B**) After transfection, Raji cells were fixed and stained with mouse anti-CD22 and anti-mouse IgG-AF647 and imaged using confocal microscopy to determine the cluster size of CD22. Data shown are average from 30 cells among 3 biological replicates; cells were analyzed using imageJ and shown as beanplots.(14) Comparisons by student’s t-test are shown relative to respective controls (****, p < 0.001; ***, p < 0.005).

### Native NEU modulate B cell activation

The CD22 receptor acts as a negative regulator of B cell activation and the organization and engagement of CD22 can alter B cell response.(50) We investigated the role of NEU1 and NEU3 in B cell activation using a calcium assay with Indo-1 dye.(51) We first asked if small molecule inhibitors of NEU enzymes had a measurable effect on B cell activation (**Figure 5**).(52) We used three different compounds: DANA, a pan-selective inhibitor of NEU enzymes; CG33300, a NEU1-selective inhibitor; and CG22600, a NEU3-selective inhibitor.(53–55) When all conditions were normalized to the control group, we observed that treating B cells with DANA increased basal activation. Additionally, cells treated with DANA and anti-IgM showed a significant increase in activation relative to control. Although these observations are consistent with native NEU activity acting as a negative regulator of B cell activation, they did not indicate which enzymes were involved. The selective NEU1 inhibitor, CG33300, showed similar effects to DANA – increased B cell activation relative to unstimulated and stimulated controls (**Figure 5B**). A selective NEU3 inhibitor, CG22600, showed similar activity – enhancing cell activation in basal and stimulated cells (**Figure 5C**). From these experiments, we concluded that native human NEU enzymes, including NEU1 and NEU3, act as negative regulators of B cell activation. We sought to confirm the role of NEU1 and NEU3 on B cell activation using siRNA knockdown conditions. Transfected B cells were subjected to Ca^2+^ assay as described above after knockdown of NEU1 or NEU3 and compared to a scRNA control (**Figure 6**). We found that both NEU1 and NEU3 knockdowns had increased basal Ca^2+^ levels in cells, consistent with our inhibitor studies.

**Figure 5.**
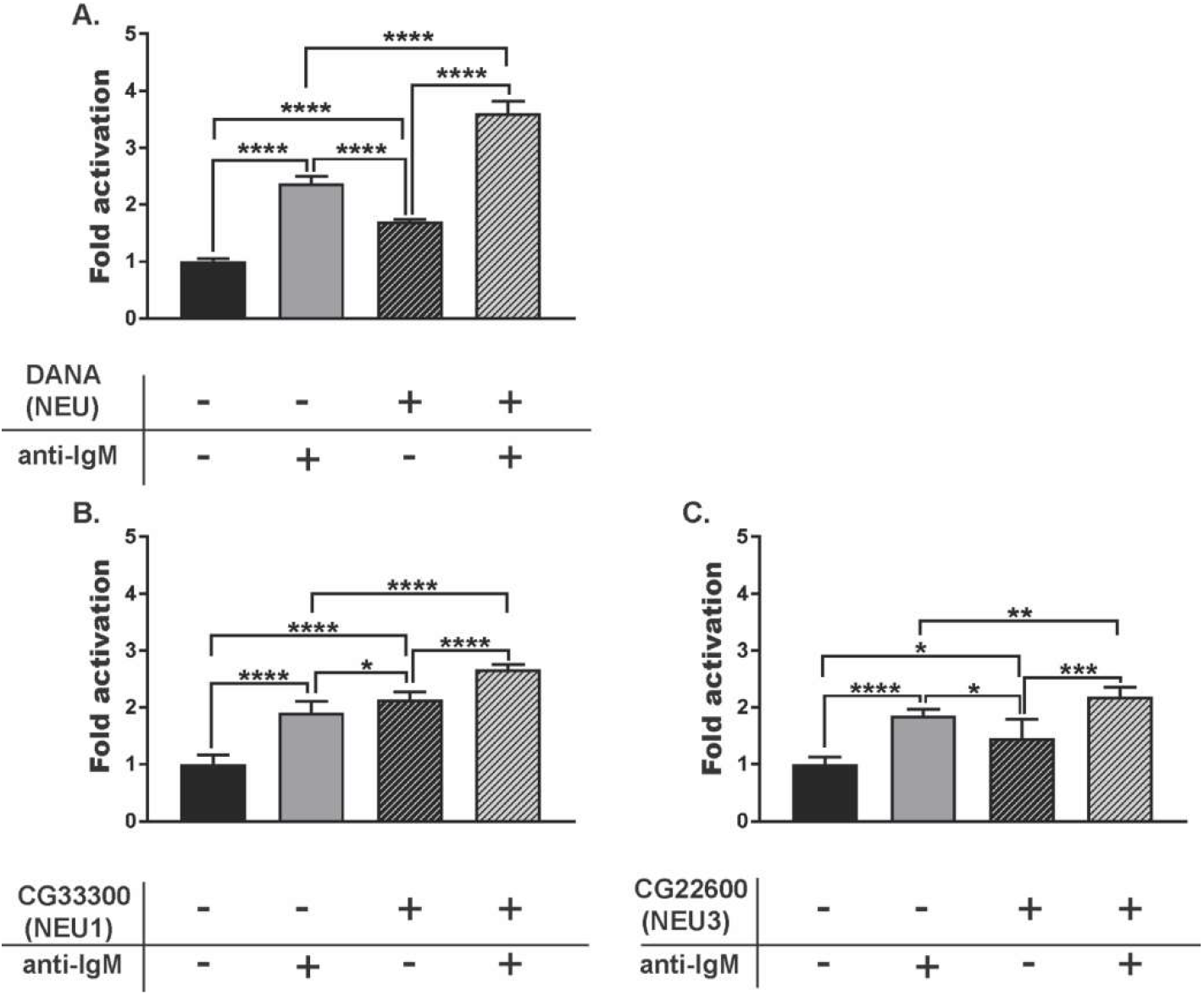
B cell response after treatment with NEU inhibitors. Raji cells were incubated at 37 °C for 30 min with NEU inhibitors: (**A**) DANA (100 µM), (**B**) CG33300, a NEU1 inhibitor (10 µM), or (**C**) CG22600, a NEU3 inhibitor (10 µM). Cells were either untreated (-, saline), or treated with inhibitor (+); followed by activation with anti-IgM. Activation of cells was monitored by observing Ca^2+^ levels by Indo-1 dye. For each treatment, 6 technical replicates from each of 3 biological replicates were performed. Responses were normalized to that of saline-treated, and unstimulated control groups and compared by student’s t-test (****, p < 0.001; ***, p < 0.005; **, p < 0.01; *, p < 0.05).

**Figure 6.**
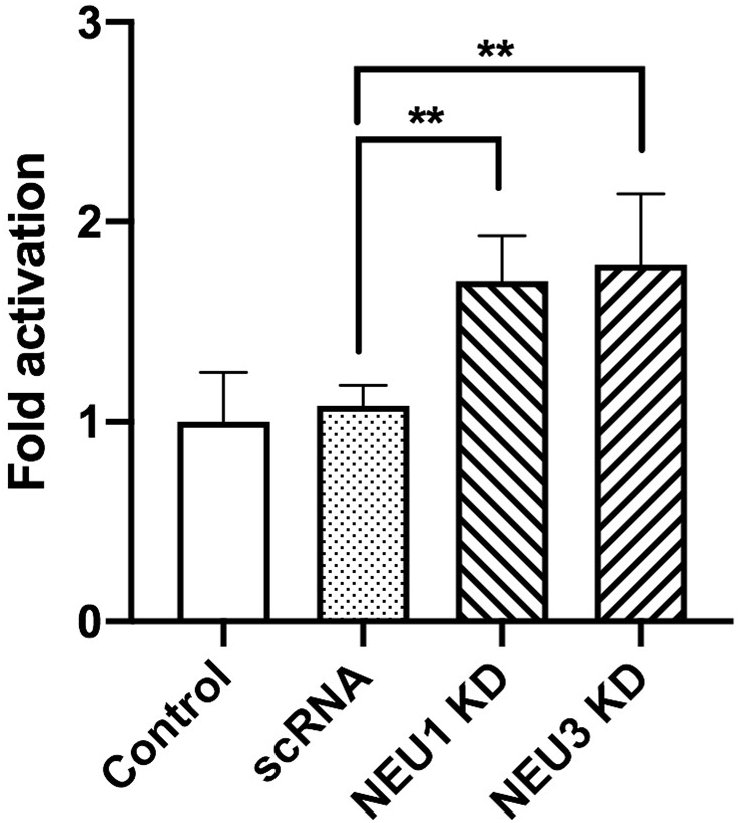
B cell calcium levels after NEU1 and NEU3 knockdown. Raji cells were transfected with siRNA targeting *Neu1, Neu3*, or a scrambled control (scRNA) using electroporation. Cells were grown for 24 h, and Ca^2+^ levels were monitored using Indo-1 dye. For each treatment, 2 technical replicates from each of 3 biological replicates were performed. Responses were normalized to that of saline-treated, and unstimulated control groups and compared by student’s t-test (**, p < 0.01).

One possible explanation for changes to B cell activation after siRNA transfection is differences in CD22 expression after treatment. We tested for changes in CD22 expression using western blot following siRNA treatments (**Figure 7, S3**). We found no significant change in CD22 expression levels in scRNA and NEU1 KD samples. However, NEU3 KD showed an unexpected decrease in CD22 expression relative to the untreated and scRNA controls. NEU3 activity has been implicated in clathrin-dependent endocytosis and could influence transport and expression of CD22.(56–59)

**Figure 7.**
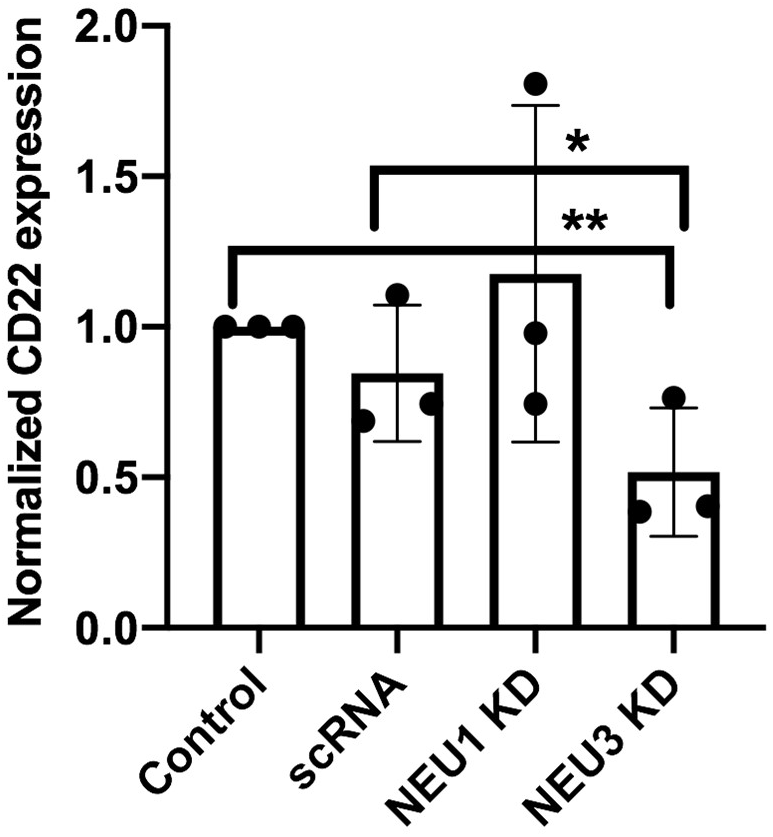
CD22 expression after NEU1 and NEU3 knockdown. Raji cells were transfected with siRNA targeting *Neu1, Neu3*, or a scrambled control (scRNA) using electroporation. Cells were allowed to grow for 24 h, and then harvested. A western blot was performed using anti-CD22 to compare expression levels and analyzed by densitometry using ImageJ (**, p<0.01; *, p<0.05).

### Exogenous NEU affect CD22 organization and B cell activation

A common strategy for probing the role of membrane sialosides in signaling is to treat cells with exogenous NEU enzymes. Typical examples include the sialidase from *Athrobacter ureafaciens* (siaAU) and NanI from *Clostridium perfringens*. These enzymes have different specificities, with siaAU having broad activity to cleave α2,3 and α2,6-linkages;(60,61) while the latter prefers α2,3-linked sialosides.(62) It is worth noting that these enzymes have different substrate specificity from human NEU isoenzymes, and may not be good biochemical proxies for the native enzymes.(63) We found that treatment of B cells with NanI and siaAU generally increased clustering of CD22 (**Figure S4A, S4C**) but not BCR (**Figure S5**). We noted that the effect on CD22 cluster size was dependent on the activity of enzyme used – with high specific activity of siaAU (10 mU/mL) reversing significant increases seen at lower activity (5 mU/mL). Analysis of B cells treated with these enzymes by SPT found that CD22 lateral mobility was significantly reduced for both NanI and siaAU treatment (**Figure 8**). This result is similar to observations with CytoD treatment, where increased clustering of CD22 was coincident with decreased lateral mobility.

**Figure 8.**
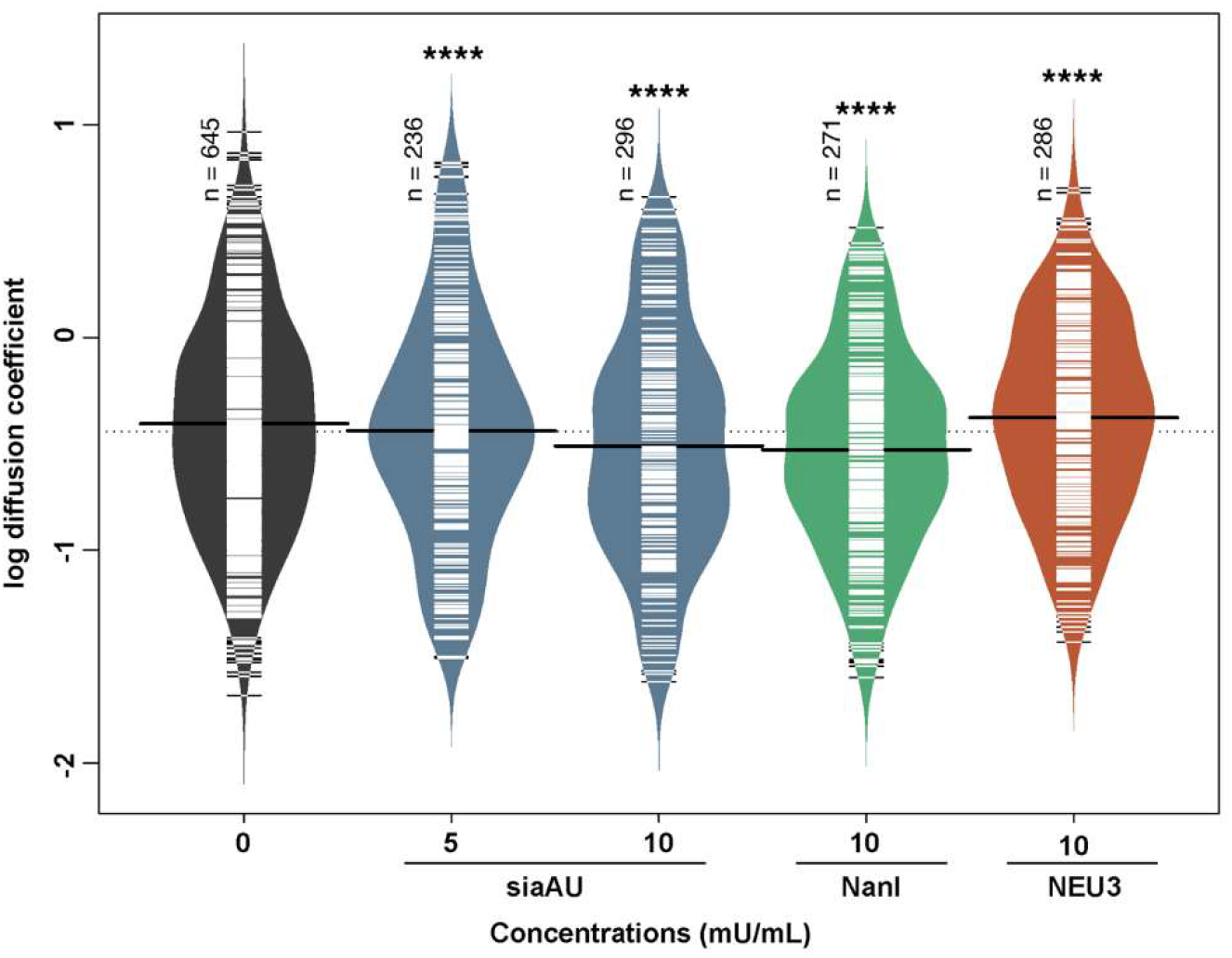
Lateral mobility of CD22 after treatment with NEU enzymes. Raji cells were treated at 37 °C for 30 min. Lateral mobility was analyzed using single-particle tracking with TIRFM videos recorded at 10 FPS for 10s.(30) Diffusion coefficients are given as log(D), where D is in units of × 10^−10^ [cm^2^s^−1^] or × 10^−2^ [μm^2^s^−1^]. Data shown are from 150 cells among 3 biological replicates, data were analyzed, and compared to control using a student’s t-test (****, p < 0.001).

We next investigated if exogenous human NEU3 enzyme had similar effects to the bacterial NEU enzymes on clustering and diffusion of CD22. When B cells were treated with NEU3 (10 mU/mL) clustering of CD22 significantly increased, consistent with the effect of the bacterial enzymes (**Figure S4B**). Measurements of the lateral mobility of CD22 after NEU3 treatment showed an increase in diffusion (**Figure 8**). We performed an analysis of B cell glycosphingolipids after exogenous NEU treatment using LC-MS (**Figure S6**),(30) in which we observed no significant changes for any of the enzyme treatments. This may suggest that changes to glycosphingolipid composition are not the major factor in changes to CD22 organization, or that these changes in composition are below the detection limit of our assay. An alternative explanation for changes to CD22 organization is that exogenous NEU enzymes modify CD22 glycosylation, thus altering homotypic clustering. Using purified CD22, we confirmed that NanI and siaAU reduced SNA staining and increased PNA staining for CD22, consistent with desialylation of the receptor in vitro (**Figure S7, S8**).

During our studies, we encountered an issue that may complicate experiments which use exogenous bacterially-produced NEU. Exogenous enzymes from bacterial sources may contain lipopolysaccharide (LPS), and this contaminant may affect lymphocyte activation(64–66) or CD22 expression.(67) We found that samples of siaAU and NanI from commercial sources contained more than 1 EU/mL (EU = endotoxin units, 1 EU = 0.1 to 0.2 ng), with final LPS concentrations in our experiments of 0.0001 to 0.01 ng/mL. We tested whether these amounts of LPS alone could affect CD22 clustering (**Figure S9**). We observed a significant increase in CD22 clustering at 0.01 ng/mL of LPS in our assay, while higher concentrations attenuated this effect (0.1 ng/mL). Additionally, determinations of B cell activation using Ca^2+^ level assays after treatment with exogenous siaAU or NanI were ambiguous in our hands (**Figure S10**). For example, siaAU at lower concentrations had similar effects to treatment with DANA (**Figure S10A** versus **Figure 5A**) despite the fact that these treatments should have opposite effects on sialic acid content on cells. Higher concentrations of siaAU attenuated this effect (**Figure S10B**), while NanI treatment showed no significant differences from control (**Figure S10C**).

## DISCUSSION

The data described here provide critical insight into the effects of native and exogenous NEU enzymes on CD22 organization on B cells. The organization of CD22 on the membrane is dependant on the lectin activity of the receptor and the availability of sialoglycoproteins in the milieu of the plasma membrane. We set out to understand if native NEU enzymes, which help regulate levels of sialyation of glycolipids and glycoproteins, could influence CD22 organization. Using confocal microscopy and single-particle tracking we demonstrated that CD22 has interactions with the cytoskeleton, though we did not resolve the nature of this interaction. We found that native NEU1 and NEU3 activity influenced both the size of CD22 clusters and their mobility within the membrane. Based on our results, we conclude that increased NEU1 activity led to smaller CD22 clusters. In contrast, increased NEU3 activity (both native or exogenous) generated larger CD22 clusters which had increased diffusion. Moreover, exogenous bacterial NEU activity generated larger CD22 clusters with decreased diffusion. These stark differences were likely the result of different substrate specificities for each enzyme. Importantly, we confirmed that LPS contamination in exogenous enzyme preparations influenced CD22 organization and could complicate attempts to use these reagents to understand the role of CD22 interactions with cis sialoside ligands. Furthermore, we confirmed that native NEU activity influenced B cell response to BCR clustering using a Ca^2+^ assay. Knockdown or chemical inhibition of both NEU1 and NEU3 enzymes resulted in increased basal activation of B cells, consistent with these enzymes acting as negative regulators of B cell stimulation. Our studies clearly support the involvement of the cytoskeleton and NEU enzymes in regulating CD22 organization and B cell activity.

Lateral mobility of immune cell receptors is complex and can be influenced by a number of factors(68) such as the lateral size of the protein,(69) cytoskeletal barriers,(70–73) the presence of membrane microdomains,(74) and crowding effects.(75) Studies have proposed CD22,(15) CD45,(76) and BCR(77,78) are associated with membrane microdomains in lymphocytes.(19,79) CD22 is not thought to have direct contacts to the cytoskeleton; though cis ligands could provide indirect contacts. In studies of CD22-cytoskeleton interactions, we found that SPT was more sensitive to changes in cluster size than confocal microscopy. Furthermore, we examined CD22-cytoskeleton interactions using a range of CytoD concentrations (2.5 - 10 µg/mL), while previous studies tested only a single concentration and found no effect.(15) The interaction of CD22 with the cytoskeleton was complex, and our data indicated that with low concentrations of CytoD (2.5 µg/mL) CD22 was found in larger clusters with reduced lateral mobility. These data cannot resolve whether CD22-cytoskeleton contacts are direct or indirect, but it is well known that this receptor is found in homotypic clusters,(5) and has cis-binding interactions with sialoglycoproteins, such as CD45.(6,15,44,80) Notably, CD45 is associated with the cytoskeleton through a spectrin-ankyrin complex which regulates its lateral mobility, providing a likely explanation for these findings.(81–83) As a Siglec that engages cis ligands, CD22 organization could be expected to be influenced by mechanisms that regulate membrane sialoglycoproteins. Previous work has found changes to CD22 organization from altered sialyltransferase expression, CD45 expression, lectin activity of CD22, and altered glycosylation sites on CD22.(15,50,84,85) This work is the first to explore the role of native NEU enzymes in CD22 organization. We found that NEU1 and NEU3 had a role in CD22 clustering, and our data suggest that isoenzymes could play disparate roles in B cell regulation. There is growing recognition that native NEU enzymes may play important roles in inflammation and immune cells.(22,30,86,87) We propose that further investigation of the role of these enzymes in B cell regulation is needed.

A common strategy to perturb CD22-ligand interactions is to treat cells with exogenous NEU. Many examples have tested the effect of exogenous bacterial NEU enzymes on CD22 organization and activity.(15,50) While these reagents may reveal aspects of CD22-ligand interactions, they may not report on the role of native NEU isoenzymes. The substrate preferences of bacterial enzymes and native NEU are different. For example, enzymes like NanI prefer glycoprotein substrates, while NEU3 prefers glycolipids.(88,89) Glycosphingolipids are a major component of membrane microdomains, and these membrane components may be modulated by NEU3 activity.(90) Bacterially-produced enzymes may also be contaminated with LPS, which we observed could alter CD22 clustering. Thus, we suggest that results based on the use of bacterially-produced enzyme be interpreted with caution; and we favor the use of small molecule inhibitors or knockdown of NEU expression to avoid this complication.

## Acknowledgements

The authors would like to thank Dr. Matthew Macauley for providing murine hybridoma cells expressing anti-CD22 antibodies. This work was supported by grants from the Natural Sciences and Engineering Research Council of Canada (NSERC RGPIN-2020-04371) and the Canadian Glycomics Network (GlycoNet).

## Author contributions

Conceived and designed experiments: CWC, HTT, CL Performed experiments and analyzed data: HTT, RC, CL Wrote the manuscript: CWC, HTT

## Conflict of interest

CWC is a co-inventor of a patent describing NEU inhibitors used in this study.

## Supplementary material

The Supplementary Material for this article can be found online at xxx.

**Figure S1. Western blots of NEU knockdowns**. Raji cells were transfected with siRNA targeting Neu1 and Neu3 using electroporation and grown for 24 hours. Western blots show the reduction of expression of hNEU1 (A) and hNEU3 (B). Shown are representative blots of three replicates for each NEU.

**Figure S2. Raji cell viability after siRNA transfection**. Raji cells were transfected with siRNA targeting Neu1, Neu3, or a scrambled control (scRNA) using electroporation and grown for 24 hours. The viability of cells from each condition was determined using trypan blue dye exclusion on a hemocytometer.

**Figure S3. Western blot of CD22 expression after NEU1 and NEU3 knockdown**. Raji cells were transfected with siRNA targeting Neu1, Neu3, or a scrambled control (scRNA) using electroporation. Cells were allowed to grow for 24 h, and then harvested. A western blot was performed using anti-CD22 to compare expression levels and analyzed by densitometry using imageJ. Shown is representative blot of three replicates.

**Figure S4. Cluster size of CD22 after treatment with NEU enzymes**. Raji cells were treated with bacterial NEU (A) or human NEU enzymes (B) at the indicated concentrations at 37 °C for 30 min. Cells were then fixed and stained with mouse anti-IgM and anti-mouse IgG-AF647 and imaged using confocal microscopy (C). Ten cells from each condition were analyzed using ImageJ and are shown as beanplots. Bottom right: confocal images of Raji cells stained with anti-CD22 antibody. Comparisons by student’s t-test are shown relative to respective controls (***, p < 0.005; *, p < 0.05).

**Figure S5. Cluster size of BCR after treatment with NEU enzymes**. Raji cells were treated with NanI, siaAU, or NEU3 enzyme at 37 °C for 30 min. Cells were then fixed and stained with mouse anti-IgM and anti-mouse IgG-AF647 and imaged using confocal microscopy. Ten cells from each condition were analyzed using ImageJ and are shown as beanplots.

**Figure S6. Glycolipid composition of Raji cells after NEU treatment**. Raji cells were treated with saline, siaAU (5 mU/mL), siaAU (10 mU/mL), NanI (10 mU/mL), or NEU3 (10 mU/mL) for 30 min at 37 °C. Cells were then subjected to glycolipid analysis using LC-MS. Data shown are the average of four replicates.

**Figure S7. Lectin blots of purified CD22 protein treated with NEU enzymes**. CD22 was purified from Raji cells using an immunoaffinity column. The protein was treated with (-) saline or (+) NEU enzymes (A) siaAU (5 mU/mL) or (B) NanI (10 mU/mL) for 30 minutes at 37 °C. Samples were then analyzed by lectin blotting with PNA, SNA, or MAL probes.

**Figure S8. Quantification of NEU-treated CD22 protein**. CD22 samples treated with (-) saline or (+) NEU enzymes (A) siaAU (5 mU/mL) or (B) NanI (10 mU/mL). Purified proteins were incubated with corresponding neuraminidase for 30 min at 37 °C. Samples were then analyzed by lectin blotting with PNA, SNA, or MAL probes, and images were quantified using densitometry.

**Figure S9. Cluster size of CD22 after treatment with LPS**. Cluster size of CD22 using confocal microscopy. Raji cells were treated with LPS at 37 °C for 30 min. Cells were then fixed and stained with mouse anti-IgM and anti-mouse IgG-AF647 and imaged using confocal microscopy. Ten cells from each condition were analyzed using ImageJ and are shown as beanplots.22 Comparisons by student’s t-test are shown relative to respective controls (*, p < 0.5).

**Figure S10. B cell response after treatment with NEU enzymes**. Raji cells were incubated at 37 °C for 30 min with NEU enzymes: (A) sialidase from Athrobacter ureafaciens (siaAU) at 5 mU/mL, (B) siaAU at 10 mU/mL, or (C) NanI at 10 mU/mL. Cells were either untreated (-, saline), or treated with enzyme (+); followed by activation with anti-IgM. Activation of cells was monitored by observing Ca2+ levels monitored by Indo-1 dye. Responses were normalized to that of saline-treated, and unstimulated control groups and compared by student’s t-test (****, p < 0.001; ***, p < 0.005; **, p < 0.01; *, p < 0.05).

