## Supporting information for "NEU1 and NEU3 enzymes alter CD22 organization on B cells"

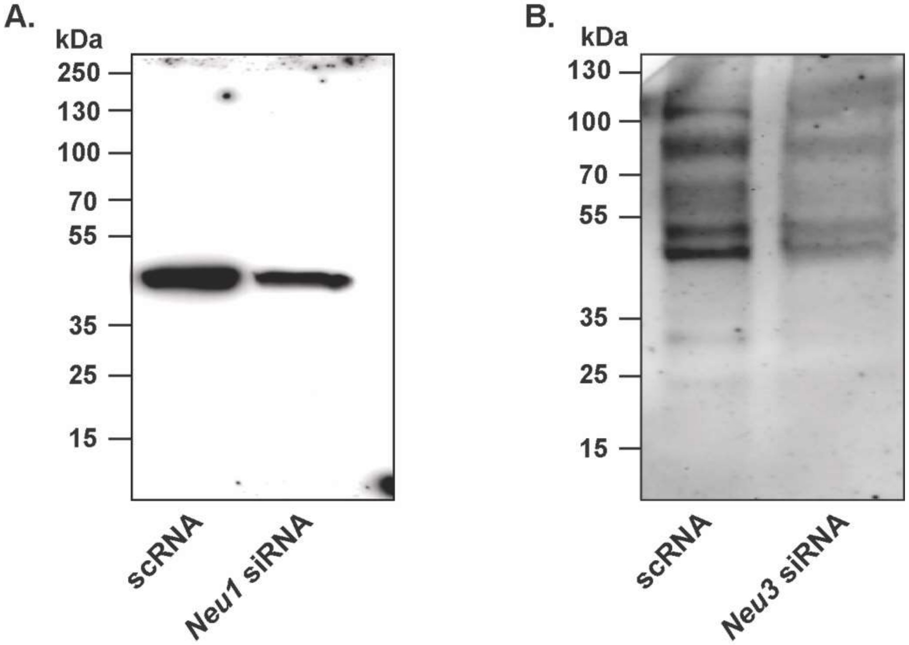

**Figure S1. Western blots of NEU knockdowns.** Raji cells were transfected with siRNA targeting *Neu1* and *Neu3* using electroporation and grown for 24 hours. Western blots show the reduction of expression of hNEU1 (**A**) and hNEU3 (**B**). Shown are representative blots of three replicates for each NEU.

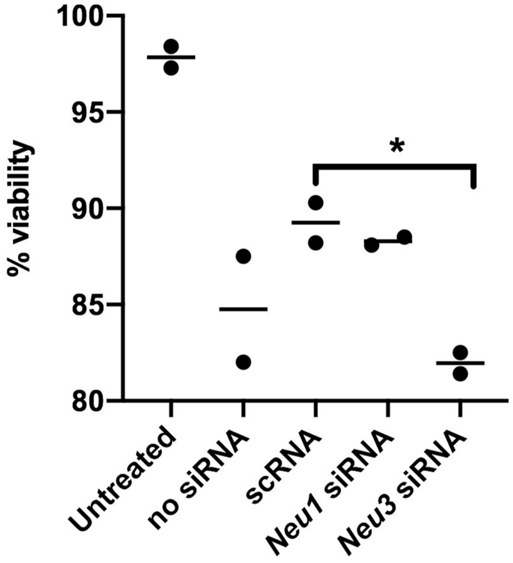

**Figure S2. Raji cell viability after siRNA transfection.** Raji cells were transfected with siRNA targeting *Neu1, Neu3,* or a scrambled control (scRNA) using electroporation and grown for 24 hours. The viability of cells from each condition was determined using trypan blue dye exclusion on a hemocytometer.

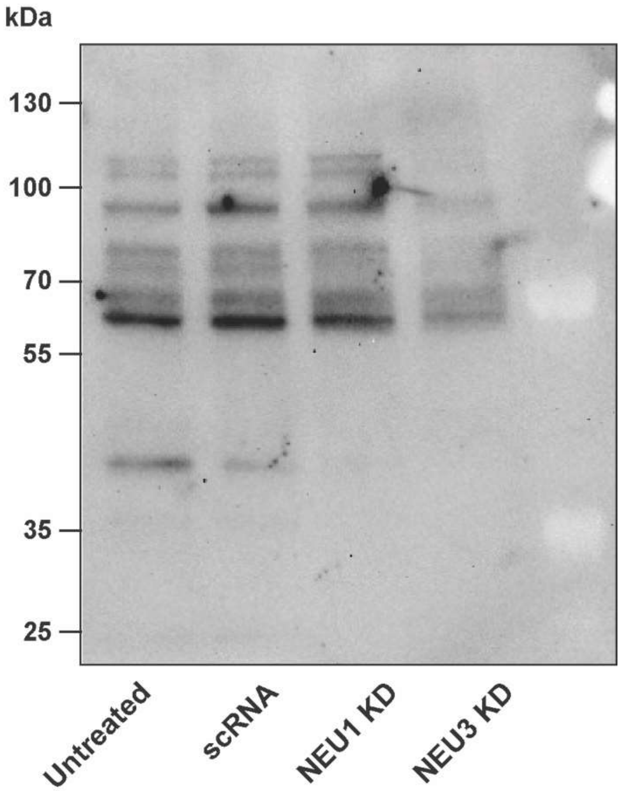

**Figure S3. Western blot of CD22 expression after NEU1 and NEU3 knockdown.** Raji cells were transfected with siRNA targeting *Neu1, Neu3*, or a scrambled control (scRNA) using electroporation. Cells were allowed to grow for 24 h, and then harvested. A western blot was performed using anti-CD22 to compare expression levels, and analyzed by densitometry using imageJ. Shown is representative blot of three replicates.

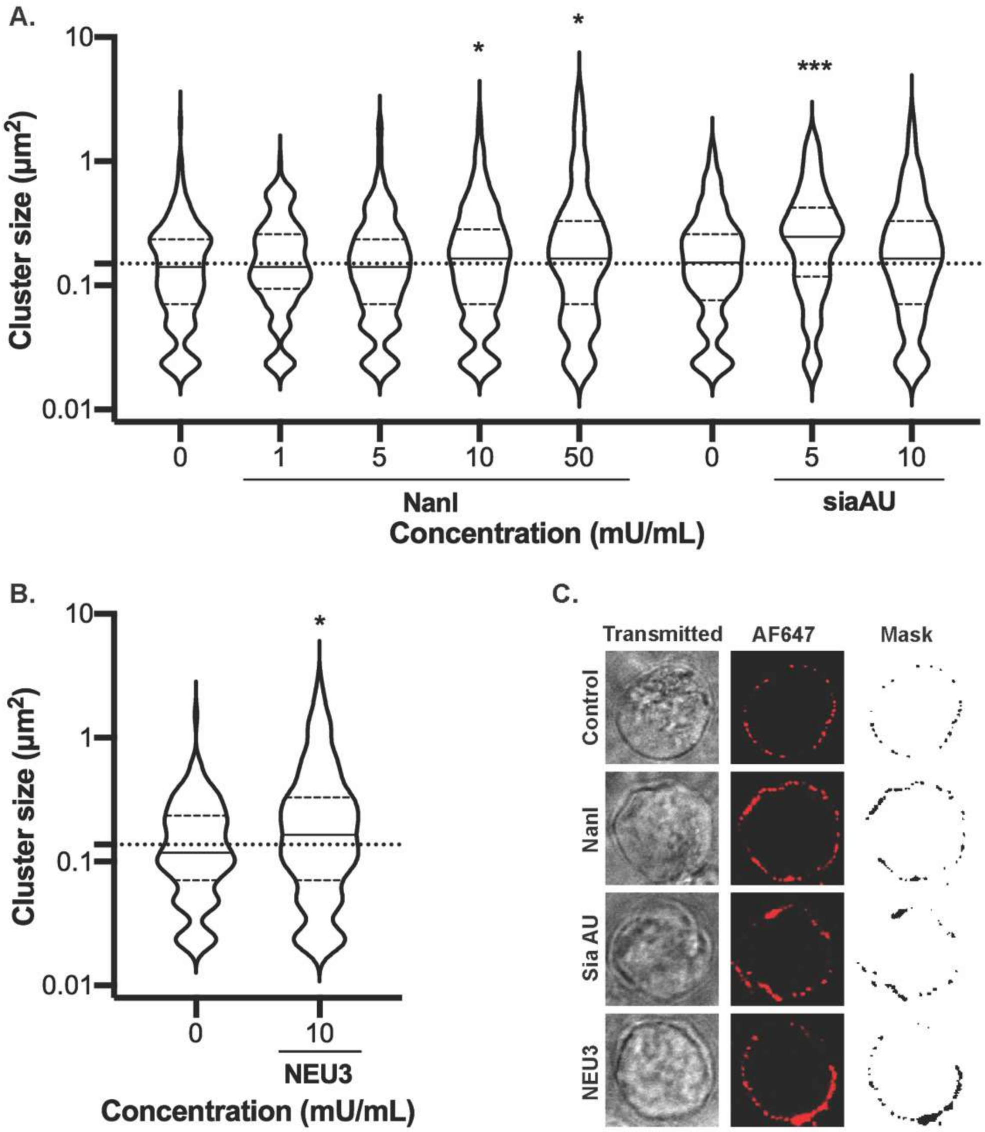

**Figure S4. Cluster size of CD22 after treatment with NEU enzymes.** Raji cells were treated with bacterial NEU (**A**) or human NEU enzymes (**B**) at the indicated concentrations at 37 °C for 30 min. Cells were then fixed and stained with mouse anti-IgM and anti-mouse IgG-AF647 and imaged using confocal microscopy (**C**). Ten cells from each condition were analyzed using ImageJ and are shown as beanplots. Bottom right: confocal images of Raji cells stained with anti-CD22 antibody. Comparisons by student's t-test are shown relative to respective controls (***, $p < 0.005$; *, $p < 0.05$).

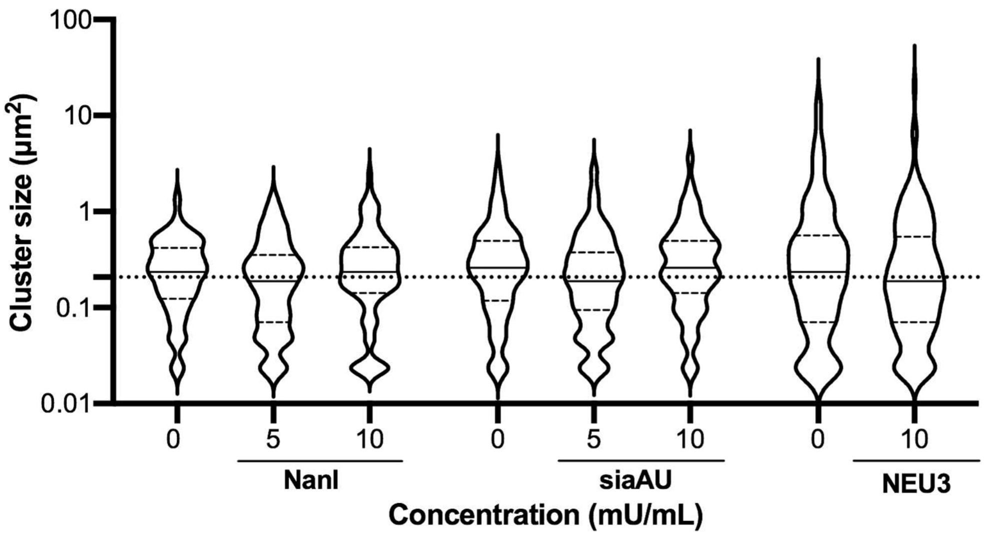

**Figure S5. Cluster size of BCR after treatment with NEU enzymes.** Raji cells were treated with NanI, siaAU, or NEU3 enzyme at 37 °C for 30 min. Cells were then fixed and stained with mouse anti-IgM and anti-mouse IgG-AF647 and imaged using confocal microscopy. Ten cells from each condition were analyzed using ImageJ and are shown as beanplots.

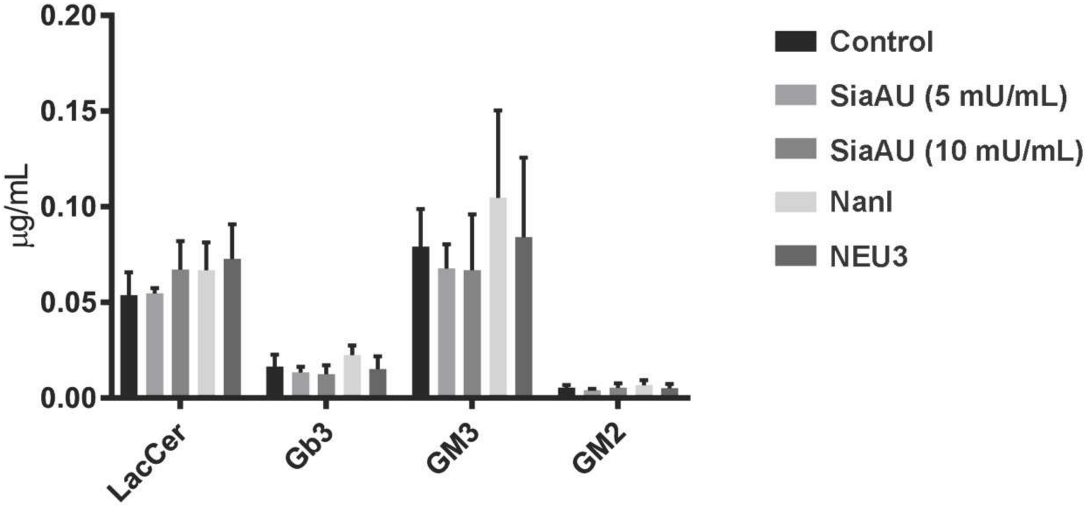

**Figure S6. Glycolipid composition of Raji cells after NEU treatment.** Raji cells were treated with saline, siaAU (5 mU/mL), siaAU (10 mU/mL), NanI (10 mU/mL), or NEU3 (10 mU/mL) for 30 min at 37 °C. Cells were then subjected to glycolipid analysis using LC-MS. Data shown are the average of four replicates.

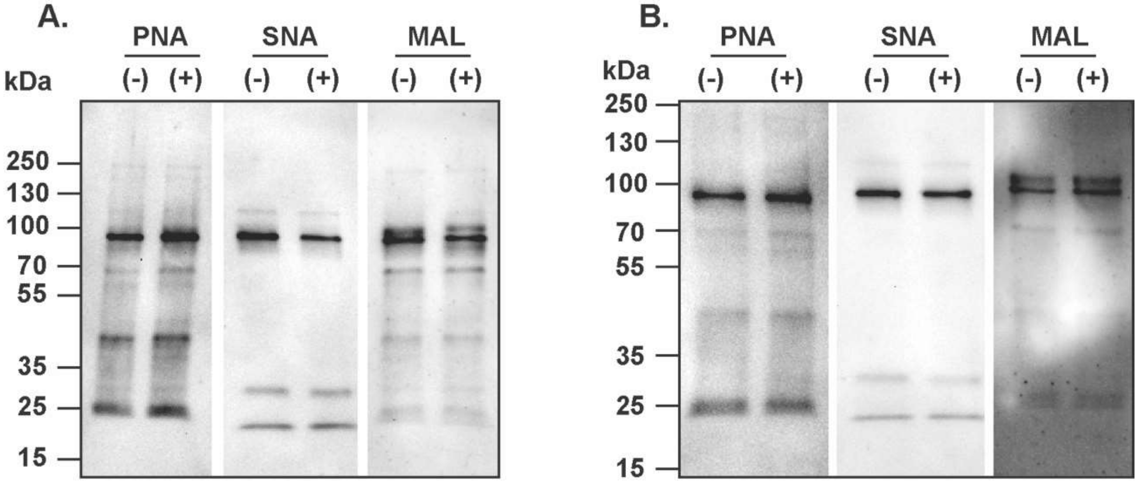

**Figure S7. Lectin blots of purified CD22 protein treated with NEU enzymes.** CD22 was purified from Raji cells using an immunoaffinity column. The protein was treated with (-) saline or (+) NEU enzymes (**A**) siaAU (5 mU/mL) or (**B**) NanI (10 mU/mL) for 30 minutes at 37 °C. Samples were then analyzed by lectin blotting with PNA, SNA, or MAL probes.

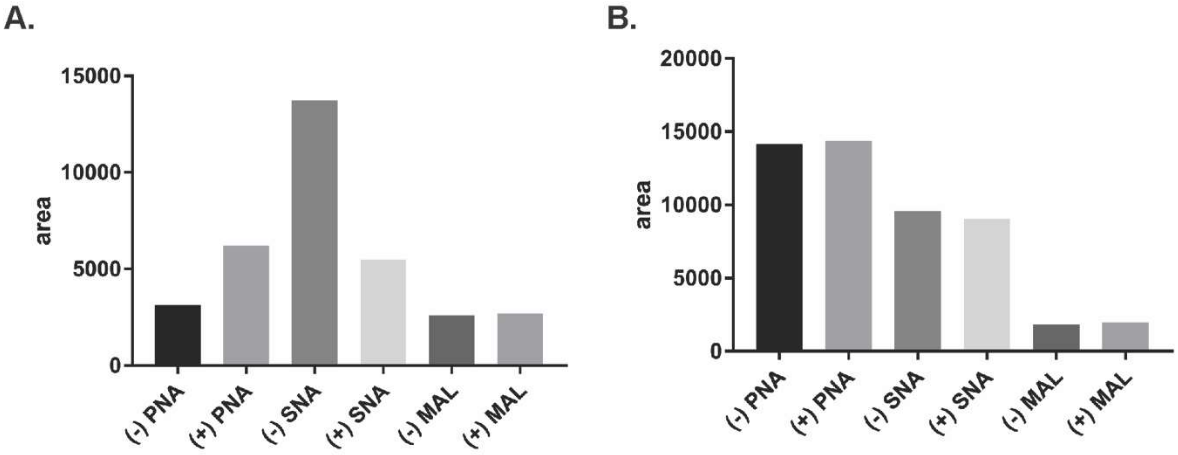

**Figure S8. Quantification of NEU-treated CD22 protein.** CD22 samples treated with (-) saline or (+) NEU enzymes (**A**) siaAU (5 mU/mL) or (**B**) NanI (10 mU/mL). Purified proteins were incubated with corresponding neuraminidase for 30 min at 37 °C. Samples were then analyzed by lectin blotting with PNA, SNA, or MAL probes, and images were quantified using densitometry.

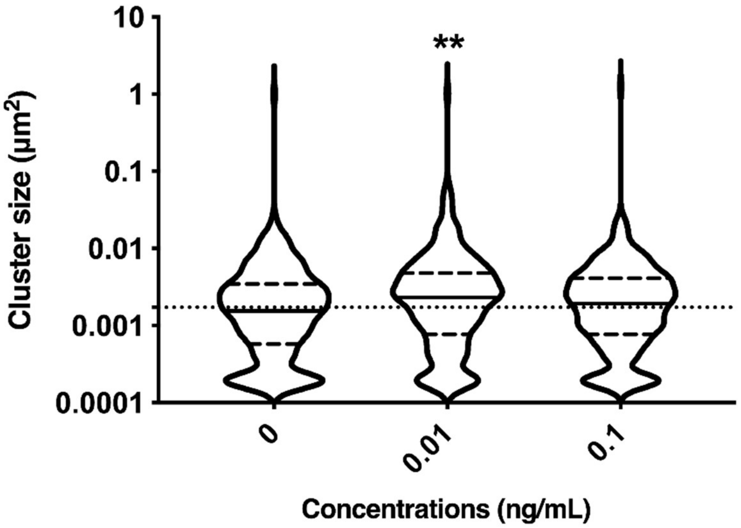

**Figure S9. Cluster size of CD22 after treatment with LPS.** Cluster size of CD22 using confocal microscopy. Raji cells were treated with LPS at 37 °C for 30 min. Cells were then fixed and stained with mouse anti-IgM and anti-mouse IgG-AF647 and imaged using confocal microscopy. Ten cells from each condition were analyzed using ImageJ and are shown as beanplots.[22] Comparisons by student's t-test are shown relative to respective controls (*, $p < 0.5$).

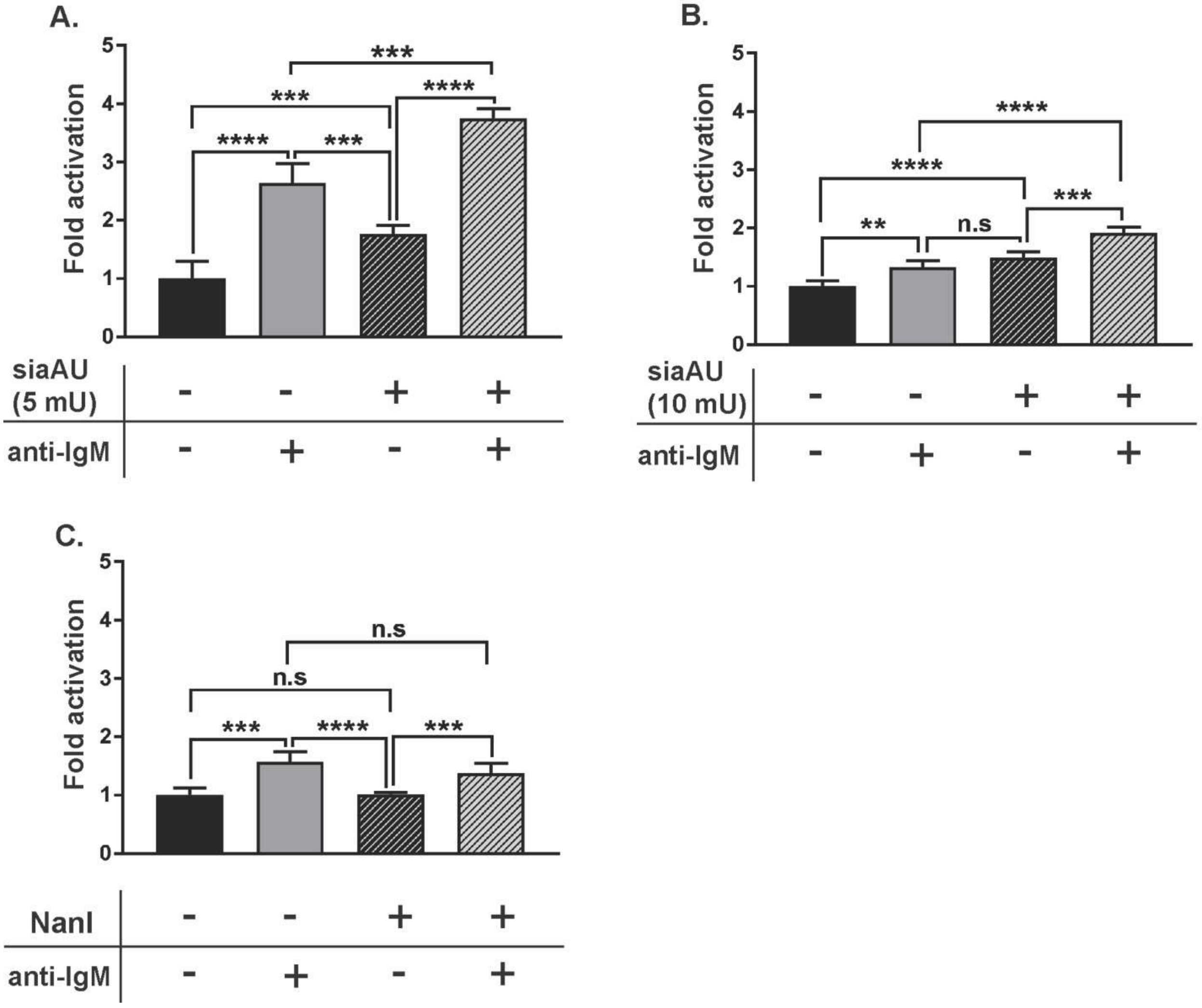

**Figure S10. B cell response after treatment with NEU enzymes.** Raji cells were incubated at 37 °C for 30 min with NEU enzymes: (**A**) sialidase from *Athrobacter ureafaciens* (siaAU) at 5 mU/mL, (**B**) siaAU at 10 mU/mL, or (**C**) NanI at 10 mU/mL. Cells were either untreated (-, saline), or treated with enzyme (+); followed by activation with anti-IgM. Activation of cells was monitored by observing $Ca^{2+}$ levels monitored by Indo-1 dye. Responses were normalized to that of saline-treated, and unstimulated control groups and compared by student's t-test (****, $p < 0.001$; ***, $p < 0.005$; **, $p < 0.01$; *, $p < 0.05$).
